# Synergy of HLA class I and II shapes the timing of antitumor immune response

**DOI:** 10.1101/2022.11.16.516740

**Authors:** Benjamin Tamás Papp, Anna Tácia Fülöp, Gergő Mihály Balogh, Balázs Koncz, Dóra Spekhardt, Máté Manczinger

**Affiliations:** Synthetic and Systems Biology Unit, Institute of Biochemistry, Biological Research Centre, Eötvös Loránd Research Network, Szeged, Hungary; HCEMM-BRC Systems Immunology Research Group, Szeged, Hungary; National Academy of Scientist Education, Szeged, Hungary; Department of Dermatology and Allergology, University of Szeged, Szeged, Hungary

## Abstract

The presentation of mutated cancer peptides to T cells by human leukocyte antigen (HLA) class I and II molecules is necessary for antitumor immune response. Both classes are diverse and the variants have distinct peptide-binding specificities. HLA class I diversity was suggested to influence antitumor immunity, however, the findings are controversial. We examined the joint effect of the two HLA classes in melanoma patients. Numerous combinations were associated with better or worse survival in metastatic melanoma patients receiving immune checkpoint blockade (ICB) immunotherapy and they also predicted the survival of ICB-naive patients. Carrying detrimental and beneficial combinations had markedly different effects in primary and metastatic samples. Detrimental combinations were associated with cytotoxic immune response in primary tumors, while metastases showed signs of immune evasion and ineffective antitumor immunity. On the contrary, beneficial combinations were associated with an active cytotoxic immune response only in metastatic samples. HLA class I and II variants in both detrimental and beneficial combinations presented melanoma-associated mutations effectively. However, detrimental combinations were more likely to present immunogenic ones. Our results provide evidence of the joint effect of HLA class I and II variants on antitumor immunity. They potentially influence the strength and timing of antitumor immune response with implications on response to therapy and patient survival.

## Introduction

The recognition of mutated cancer peptides (neopeptides) by the immune system is essential for destroying cancer^1^. Neopeptides are presented by HLA molecules to T cells, which is the prerequisite for the immune recognition of tumor cells^1^. HLA molecules can be classified into two classes^2^. HLA class I (HLA-I) molecules are expressed by all nucleated cells and present short, usually nine amino acid long peptides generated inside the cell^2^. HLA class II (HLA-II) molecules are expressed on the surface of professional antigen-presenting cells and a relevant fraction of cancer cells, and usually present 15 amino acid long peptides^2,3^.

The HLA-I-dependent presentation of neopeptides can induce a cytotoxic, CD8+ T cell-mediated immune response, while their presentation by HLA-II molecules can activate CD4+ T cells^2,4^. The activation of CD4+ T cells is needed for the augmentation of cytotoxic immune response and for the development of humoral immune response^2,4^. While the importance of HLA-I presented neopeptides has been extensively researched in the previous two decades^5–7^, the relevance of HLA-II molecules in antitumor immunity emerged only recently^3,8,9^. It was reported that vaccination with MHC-II neoantigens induces a robust cytotoxic antitumoral immune response in a murine model^9^. In another study, MHC-I neoantigens alone were not able to induce successful tumor destruction, while their co-occurrence with MHC-II neoantigens resulted in tumor clearance^8^. Moreover, it was suggested that the effective HLA-II presentation of driver mutations is a significant negative selection pressure on their occurrence^10^. In line with these findings, both HLA-I and HLA-II neoantigen burden predicts the survival of cancer immunotherapy patients^11,12^.

Both HLA classes are highly variable and thousands of alleles have already been registered with varying specificities^13^. The variability in HLA-I alleles has been shown to affect antitumor immunity: homozygosity, certain supertypes and evolutionary divergence of HLA-I molecules were suggested to influence the response to cancer immunotherapy^6,14^. However, subsequent studies questioned the robustness of their predictive power^15,16^. Notely, we have previously shown that the peptide-binding repertoire size of HLA-I variants also affects the efficacy of antitumor immune response^7^.

As the presentation of neopeptides by both HLA classes is essential for antitumor immunity, we aimed to examine the joint effects of their variants in melanoma patients. We hypothesized that the similarities and differences between their peptide-binding influence antitumor immune response. Accordingly, the limited robustness of the reported findings for HLA-I diversity could be explained by the different HLA-II alleles carried by patients. We identified numerous combinations with remarkable effects on survival. These effects were much stronger and more robust than for the two classes separately. Surprisingly, detrimental combinations dominated our findings, which were able to present melanoma-associated neopeptides with considerable efficiency. They induced intensive inflammation already in primary tumors leading to immune evasion in metastases. At the same time, beneficial combinations were associated with a cytotoxic immune response in metastases only. Our findings suggest a strong allele-specific interplay between the recognition of neopeptides by HLA-I and II molecules. The study not only provides accurate biomarkers for clinical use but also highlights the relevance of the optimal timing of immunotherapy.

## Results

### HLA class I and II alleles predict the survival of ICB-treated and ICB-naive melanoma patients in interaction

To examine the joint effect of HLA class I and II molecules, we first collected pairs of HLA-I and II molecules carried by melanoma patients in a large ICB cohort^6^. The cohort contained 269 melanoma patients treated either with CTLA-4 or PD-1 inhibitors. Similarly to previous studies^5,10^, we first generated all possible HLA molecules for each patient as follows. In the case of HLA class I molecules and the HLA class II molecule HLA-DRB1, one of the two chains (β2-microglobulin and HLA-DRA) is invariable^2,10^. Therefore, we took only alleles decoding the variable chain into account. At the same time, both α and β chains of HLA-DP and HLA-DQ molecules are variable and influence peptide-binding specificity^10^. We generated all possible variations between the α and the β chains and treated each variation as a separate HLA-DP or DQ molecule in the subsequent analyses. The most common combination contained the HLA class I A*02:01 and the class II DPA1*01:03-DPB1*04:01 molecules and was carried by 28% percent of patients. It was followed by the pairs of A*01:01 and DPA1*01:03-DPB1*04:01 and C*07:01 and DPA1*01:03-DPB*04:01 carried by 21% and 19% of patients, respectively (**Table S1**, **Figure S1**).

Next, we constructed Cox survival models with interaction terms:

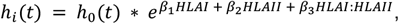

where *h_i_*(*t*) is the hazard function of individual *i, h*_0_(*t*) is the baseline hazard function; *HLAI* and *HLAII* are logical covariates indicating the presence of a given HLA-I and HLA-II molecule, respectively; *HLAI: HLAII* is the interaction term; *β*_1_, *β*_2_ and *β*_3_ are regression coefficients of the two logical covariates and the interaction term, respectively. A significant interaction term suggests that the effect of one allele in the model is influenced by the presence of another one. We found 11 combinations having significant interaction terms in these models. As HLA genes are in strong linkage disequilibrium (LD), it is possible that certain interaction effects are not directly caused by the alleles involved in the models. To this end, we controlled for the presence of other combinations in Cox models and discarded the B*08:01-DPA1*01:03-DPB1*04:01 (see Methods for details). Next, we focused on the remaining ten combinations (**Figure 1A, Table S2**.). Surprisingly, the coefficients were positive (i.e. the hazard ratio values were above one) in 8 models indicating that the HLA-I and II molecules have a negative joint effect on survival compared to when they occur alone. The coefficients of the interaction terms were negative (i.e. the hazard ratio values were below one) in two models suggesting that the two molecules together have a positive effect on survival compared to their sole presence.

**Figure 1.**
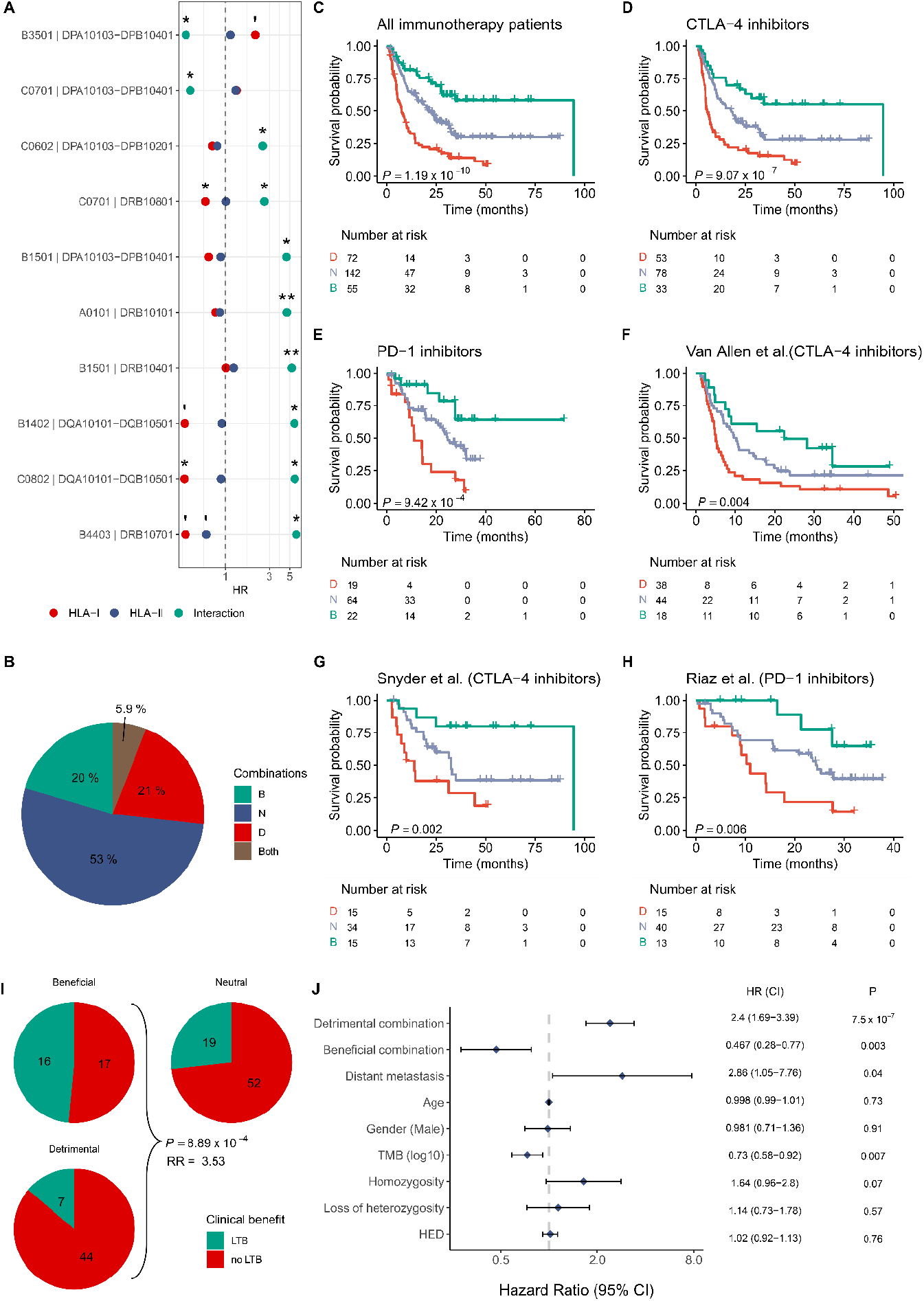
HLA-I and II alleles predict the survival of ICB-treated melanoma patients in interaction. **A)** The hazard ratio (HR) values associated with HLA-I and II molecules and their interaction in Cox models. Two-sided P values of z-statistics < 0.005 (**), < 0.05 (*), < 0.1 (‘) are highlighted. **B)** The pie chart shows the fraction of individuals carrying beneficial (B) or detrimental (D) combinations, both of them or only neutral ones (N). **C–H)** Overall survival of cancer immunotherapy patients carrying D, B or only N combinations. Two-sided log-rank test P values are shown. Vertical axes indicate the probability of patient survival. **I,** The pie chart shows the fraction of patients in the D, N and B combination groups having or not having long-term benefit (LTB) after immunotherapy. Exact patient counts in each subgroup are indicated. The two-sided P value of a Fisher’s exact test and relative risk (RR) value are indicated. **J,** Summary of a multiple Cox model, which was made using the whole immunotherapy cohort (n = 269). The effects of B and D combinations were compared to N ones in the model. Blue squares indicate hazard ratios and horizontal lines indicate 95% confidence intervals. Two-sided *P* values of *z*-statistics are shown. The two-sided log-rank test P value of the model was 3.7*10^-10^. TMB: tumor mutational burden, HED: HLA evolutionary divergence

A prominent example of the first type of combinations is HLA-B*15:01 and DRB1*04:01. The two molecules had no effect on survival alone but they had a devastating effect together (**Figure 1A, Table S2, Figure S2**). The negative effect of HLA-B*15:01 on survival was already reported^6^. Our result suggests that it is not directly related to the HLA-I allele, but its joint effect with HLA-DRB1*04:01. The second pattern is well illustrated by the HLA-C*07:01-DPA1*01:03-DPB1*04:01 combination. While the two molecules had neutral effects alone, they improved survival in combination (**Figure 1A, Table S2, Figure S2**). Interestingly, HLA-C*07:01 and HLA-DPA1*01:03-DPB1*04:01 were found in both beneficial and detrimental combinations, suggesting that the effect of a given allele can be markedly different when it is in combination with different alleles.

We also examined the effect of all HLA-I and HLA-II molecules separately in Cox models (**Table S3**). Only one HLA-I allele, HLA-B*15:01, and one HLA-II allele, HLA-DRB1*04:01 had a significant effect on survival. As shown, these molecules exerted their negative effect in combination only (**Figure 1A, Table S2, Figure S2**). Accordingly, we conclude that the findings for separate HLA-I and II alleles are only artifacts caused by the joint effect of the two classes.

Next, we examined the survival of patients carrying any of the beneficial and/or detrimental combinations in the whole cohort (**Figure 1C**) and its sub-cohorts. Importantly, 45% of patients carried at least one combination (**Figure 1B**). The carriage of detrimental and/or beneficial combinations robustly predicted the survival of both aCTLA-4 (**Figure 1D, F, G, Figure S3**) and aPD1-treated patients (**Figure 1E, H, Figure S3**). Surprisingly, patients carrying both beneficial and detrimental combinations had similar survival to patients carrying only detrimental ones (**Figure S3**), which suggests that the effect of detrimental combinations is superior to beneficial ones. According to these results, we classified individuals carrying both combination types into the detrimental group in further analyses. As expected, beneficial combinations were associated with a 3.5-times higher chance of responding to ICB therapy than detrimental ones (**Figure 1I**). Finally, the effect of combinations on survival remained especially strong in a multiple cox model involving demographic factors, tumor mutational burden and other HLA-associated features as covariates (**Figure 1J**). Importantly, the carriage of detrimental or beneficial combinations was the strongest determinant of patient survival.

In sum, we found a strong predictive power of beneficial and detrimental allele combinations on the survival of ICB-treated patients. Next, we examined whether these effects are prominent only in the ICB setting or can also be detected in ICB-naive patients of the TCGA database. The combinations predicted the overall survival of patients diagnosed with metastases (**Figure 2B**). On the contrary, they did not have any effect on the survival of patients, whose diagnosis was made in localized disease stage (stage I or stage II) (**Figure 2A**). To get a more detailed picture of the disease history of each patient, we created special plots of TCGA patients, on which relapse, disease outcome and the time spent in each stage can be followed (**Figure 2C**). Surprisingly, the chance for locoregional recurrence was higher in the beneficial combination group but only during the primary, not the metastatic stage (**Figure 2D**). Carrying detrimental combinations was associated with a 2.9-times lower survival rate than carrying beneficial ones, if the diagnosis was made in a metastatic stage, while this did not stand for patients diagnosed with primary tumors (**Figure 2E**).

**Figure 2.**
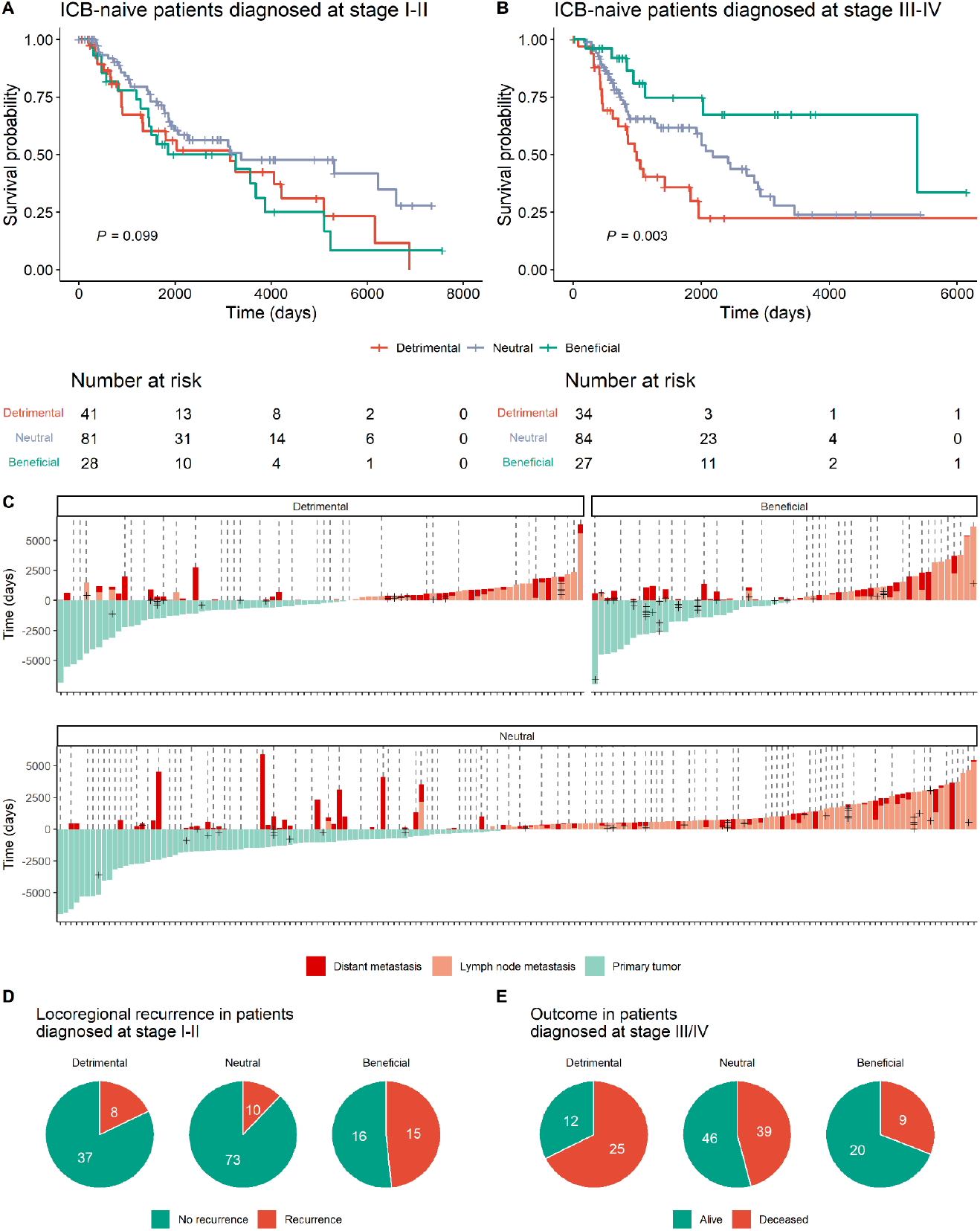
Combinations of HLA class I and II molecules determine the survival of ICB-naive melanoma patients. **A-B)** The Kaplan-Meier curves indicate the overall survival of patients carrying different HLA allele combinations and diagnosed at primary (**A**) and metastatic (**B**) stages. Log-rank test P-values are indicated. **C)** The plots indicate the medical histories of patients belonging to different HLA combination groups. Zero on the vertical axis indicates the first progression event, the time spent at the primary stage is marked in the negative range. Different colors correspond to different stages. The crosses represent locoregional recurrence. A continuous dashed line indicates that the patient was alive at the time of the last observation. **D)** The frequency of locoregional recurrence when diagnosed at stage I and II. Locoregional recurrence was 3.74-times more common in patients carrying beneficial combinations compared to others (Fisher’s exact test P: 9.45×10^-5^; for stage III-IV patients, Fisher’s exact test P: 1). **F)** The distribution of disease outcomes in different HLA combination groups of melanoma patients diagnosed at stage III/IV. Patients carrying detrimental combinations were 1.55-times less likely to survive compared with others (Fisher’s exact test P: 0.008; for patients diagnosed with primary tumors, Fisher’s exact test P: 0.49). On panels D and E, the numbers indicate patient counts in each group.

### Diametrically opposed effect of detrimental and beneficial allele combinations in primary and metastatic tumors

The results of the survival analysis indicated markedly different effects of beneficial and detrimental HLA combinations on the survival of patients diagnosed with stage I-II and stage III-IV disease. To this end, we examined the tumor immune microenvironment separately in primary and metastatic melanoma samples in the TCGA database. We used data from previous studies^17^ and also carried out immune deconvolution using state-of-the-art methods (see Methods for details).

In primary samples, detrimental combinations were associated with elevated lymphocyte and NK cell count in the tumor microenvironment (**Fig 3, A and B**). Lymphocytes were mainly B cells and CD8+ T lymphocytes, but not CD4+ T lymphocytes (**Fig 3, C to E**). A higher cytotoxicity score (**Fig 3F**) and increased Shannon entropy of TCRs (**Fig 3G**) were also detected in these samples. The findings suggest that detrimental combinations cause a cytotoxic immune response in primary tumors. On the contrary, beneficial combinations were associated with an especially low lymphocyte, NK and B cell count, cytotoxicity score and TCR diversity (**Fig 3, A to G**). Interestingly, the number of activated dendritic cells was elevated in these samples (**Fig 3, H and I**) suggesting active antigen presentation in these samples^18^. We examined the distribution of tumor microenvironment subtypes according to Bagaev et al.^19^. As expected, samples in the detrimental combination group were more likely to have an inflamed phenotype (**Fig 3J,** Fisher’s exact test P: 0.01, relative risk: 5.6 vs. other primary tumors).

**Figure 3.**
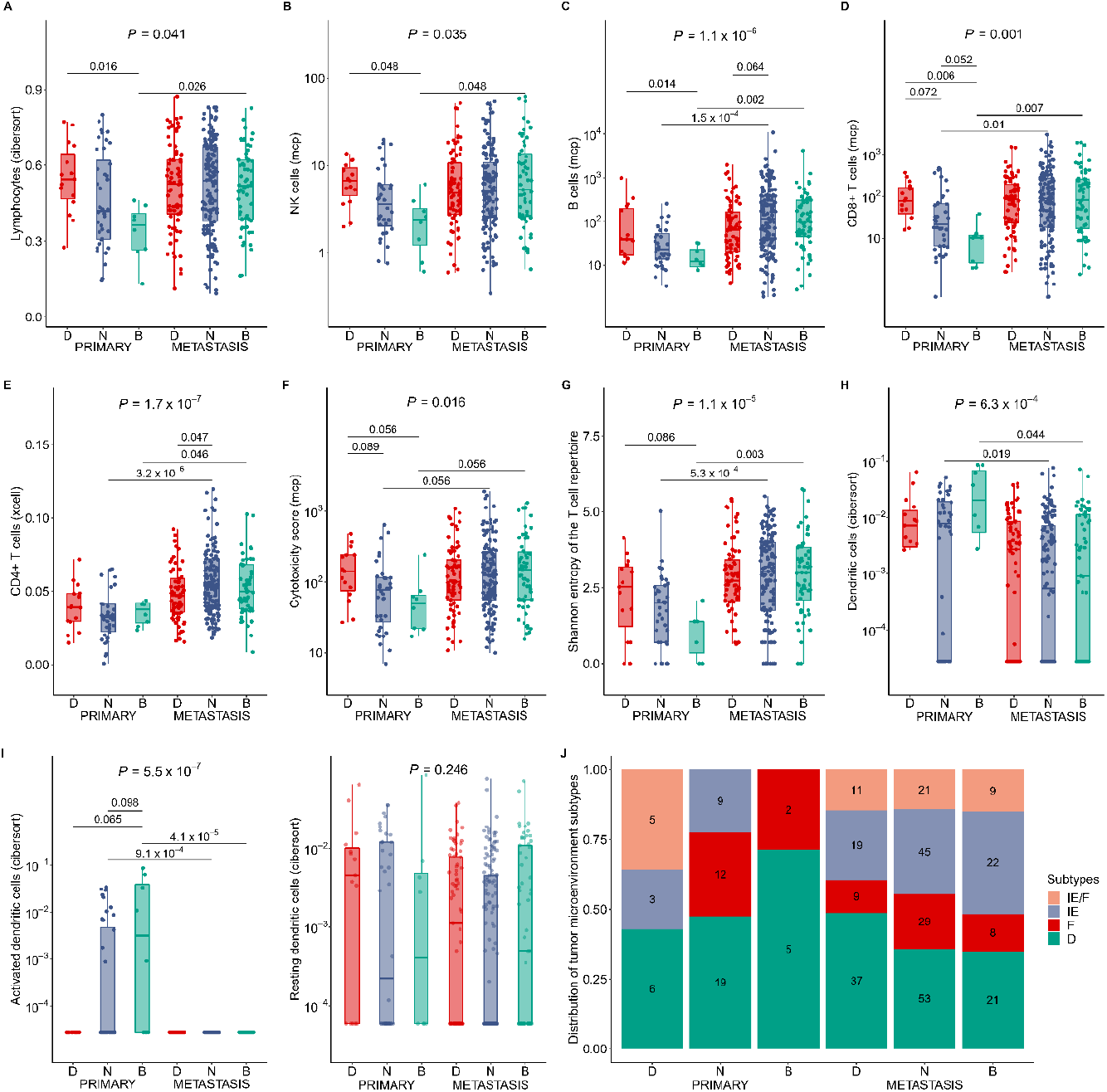
The immune microenvironment of melanoma samples. **A)** The relative fraction of lymphocytes in samples as reported previously^17^ (n = 15, 40 and 8 primary tumor samples carrying D, N and B combinations, respectively. n = 77, 150 and 61 metastatic tumor samples carrying D, N and B combinations, respectively). **B-D)** The abundance of NK cells, B cells and CD8+ T cells determined with the MCP-counter algorithm^20^ (n=15, 36 and 8 primary tumor samples carrying D, N and B combinations, respectively. n = 77, 150 and 61 metastatic tumor samples carrying D, N and B combinations, respectively). **E)** The abundance of CD4+ T cells determined with the xCell algorithm^21^ (n = 15, 36 and 8 primary tumor samples carrying D, N and B combinations, respectively. n = 77, 150 and 61 metastatic tumor samples carrying D, N and B combinations, respectively). **F)** The cytotoxicity scores in samples as calculated by the MCP-counter algorithm (sample counts are the same as for panels B to D). **G)** The Shannon entropy of the T cells in tumor samples repertoire as reported by ref.^17^ (n = 15, 37 and 7 primary tumor samples carrying D, N and B combinations, respectively. n = 74, 139 and 56 metastatic tumor samples carrying D, N and B combinations, respectively). **H-I)** The relative fraction of total, activated and resting dendritic cells in samples as reported previously^17^ (sample counts are the same as for panel G). **J)** The distribution of subtypes of the tumor microenvironment (IE/F - immune-enriched, fibrotic; IE - immune-enriched, non-fibrotic; F - fibrotic; and D - immune-depleted) in melanoma samples. The numbers indicate sample counts belonging to different subtypes. On panels A to I, P-values of Kruskal-Wallis tests are indicated (on top). On the lines above the boxplots, FDR-corrected P-values of pairwise Wilcoxon’s rank-sum tests are indicated. Only P-values less than 0.1 are shown for visualization purposes. The bottom and top of the boxes indicate the first and third quartile, horizontal lines indicate the median, and vertical lines indicate the first quartile - l.5*IQR and the third quartile + 1.5*IQR.

We expected to find prominent immune infiltration in metastatic samples compared to primary ones as these samples contain a higher number of mutations. Indeed, we found elevated NK, B, CD4+ and CD8+ T cell counts, elevated cytotoxicity score and TCR diversity in metastases (**Fig. S4**). However, when we compared primary and metastatic samples belonging to each combination group, the picture changed: in the detrimental combination group, there was no significant difference in any cell type (**Fig 3, A to G**). At the same time, in the beneficial combination group, the levels of CD4+ and CD8+ lymphocytes, B cells, NK cells and cytotoxicity scores were much higher in metastatic than in primary samples (**Fig 3, A to G**). We found similar, but less pronounced trends in samples having only neutral combinations. The findings suggest the presence of an active cytotoxic antitumor immune response in primary samples of the detrimental group, and that the tumor immune microenvironment does not significantly change in metastases. On the other hand, primary samples in the beneficial group showed a lack of cytotoxic antitumor immune response and the dominance of dendritic cells, while in metastatic samples we found a prominent cytotoxic immune response.

### Detrimental HLA class I and II combinations are associated with prominent immune evasion in metastases

The results so far indicated that both detrimental and beneficial allele combinations are potentially associated with an effective cytotoxic immune response in certain stages of disease progression. However, the poor response to ICB therapy and the unfavorable survival of patients in the detrimental combination group suggested that these HLA class I and II combinations potentially promote immune evasion during cancer progression.

To test this assumption, we first aimed to determine the strength of immunological selection on primary tumor samples in each combination group. It is reported that antitumor immunity restricts the genetic diversity of tumors resulting in low intratumor heterogeneity (ITH)^22^. We downloaded ITH data generated by the CloneSig software^23^ and found that primary tumors in the detrimental combination group are formed by an especially low number of clones (**Fig 4A**) and the maximal distance between mutational signatures of two clones in the same sample tended to be lower (**Fig 4B**) reflecting strictly constrained evolutionary trajectories of cancer cell clones. Considering the immune phenotype of samples belonging to this group, low ITH is most likely caused by immune selection pressure. Samples with beneficial combinations showed the opposite trend: an especially high number of clones were present and their mutational signatures were more dissimilar suggesting the lack of immunological selection (**Fig 4, A and B**). In line with expectation, metastatic samples of the detrimental combination group had elevated ITH reaching a level similar to samples in other combination groups (**Fig 4, A and B**), which suggests the release of immunological selection in these samples.

**Figure 4.**
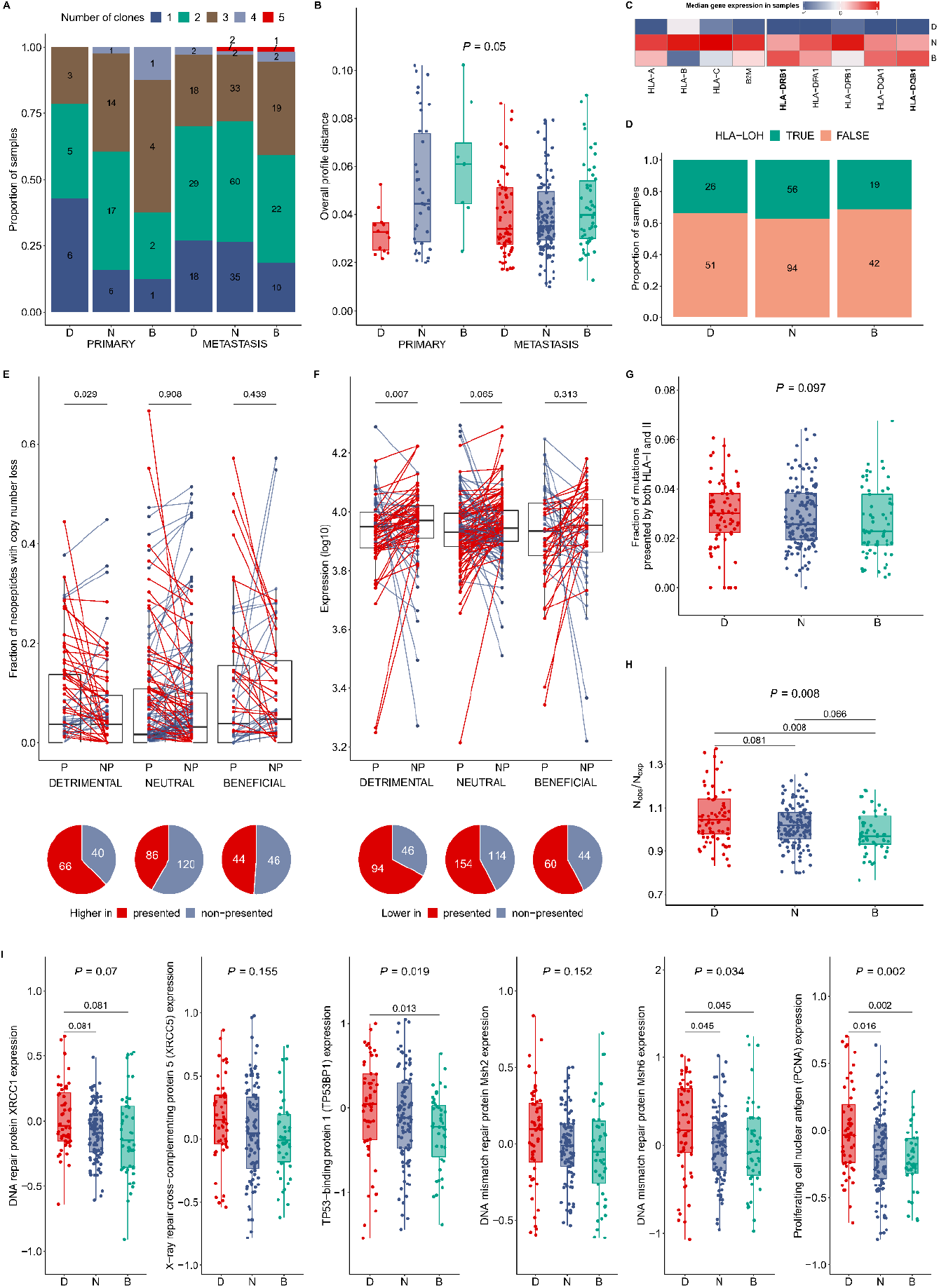
Signs of immune evasion in metastatic samples. **A)** The proportion of samples containing a given number of clones is shown in each group. The numbers indicate exact sample counts. Primary samples carrying beneficial combinations were 2.4-times more likely to consist of only one clone than other primary samples (one-sided P value of Fisher’s exact test: 0.038) **B)** The maximal distance between the mutational signatures of any two clones in a given sample (n = 14, 38 and 8 primary tumor samples carrying D, N and B combinations, respectively. n = 67, 132 and 54 metastatic tumor samples carrying D, N and B combinations, respectively). **C)** Median mRNA expression of HLA molecules in metastatic sample groups (n = 77, 150 and 61 samples in D, N and B groups). Expression values were re-scaled in each column using Z-score transformation. Bold characters indicate HLA genes showing significantly different expression in different groups (i.e. Kruskal-Wallis test P < 0.05). **D)** The proportion of samples with loss of heterozygosity (LOH) events at HLA-I, HLA-II or β2M loci. **E)** On paired boxplots, the fraction of presented (P) and non-presented (NP) subclonal mutations that are associated with copy number loss (CNL) is shown in each metastatic sample (n = 77, 149 and 60 samples in the D, N and B combination group). P-values of one-sided paired Wilcoxon’s rank-sum tests are indicated. The pie charts show the number of samples in each combination group, in which CNL is more common in presented than in non-presented peptides (RR: 2.11 in the detrimental group vs. other groups, two-sided Fisher’s exact test P: 0.005). **F)** On paired boxplots, the expression of clonal presented (P) and non-presented (NP) neopeptides without copy number alteration (CNA) is shown in metastatic samples (n = 70, 134 and 52 samples carrying D, N and B combinations). On the lines, P-values of one-sided paired Wilcoxon’s rank-sum tests are indicated. Pie charts indicate the number of samples, in which the expression of genes encoding presented neopeptides is lower than of the ones encoding non-presented neopeptides (RR: 1.51 in the detrimental group vs. other groups, two-sided Fisher’s exact test P: 0.054). **G)** The fraction of mutations presented strongly by both HLA-I and HLA-II alleles is shown for each sample in the three combination groups (n = 77, 149 and 61 in the D, N and B groups, respectively). **H)** The ratio of the observed and the expected number of missense mutations in sample groups (n = 77, 149 and 61 in the D, N and B groups, respectively). **I)** Expression of DNA repair proteins in different sample groups (n = 56, 105 and 46 in the D, N and B groups, respectively). On the top of panels B, G, H and I, the P-values of Kruskal-Wallis tests are indicated. In panels H and I, FDR-corrected P-values of pairwise Wilcoxon’s rank-sum tests are indicated on segments. Only values lower than 0.1 are shown for visualization purposes.

Next, we examined the signs of immune evasion in metastatic samples. Tumors frequently downregulate the expression or completely lose one or both copies of genes that encode proteins involved in antigen presentation, like HLA molecules or β2-microglobulin^24,25^. As expected, the expression of these genes, especially HLA-DQB1 and DRB1 was lower in the detrimental combination group than in other groups (**Fig 4C**). At the same time, there was no significant alteration in their copy numbers (**Fig 4D**). Similarly, tumors frequently downregulate or lose neoantigen-coding genes to avoid immune recognition^26^. We examined the expression and copy number alteration of genes encoding HLA-presented neopeptides in metastatic samples (see Methods for details). As controls, we selected neopeptides that are not presented by HLA molecules. In samples of the detrimental combination group, the loss of one or both copies of genes encoding HLA-presented mutations was more common than the ones encoding non-HLA-presented mutations (**Fig 4E**). Similarly, we found that genes encoding HLA-presented mutations have lower expression than the ones encoding non-HLA-presented mutations (**Fig 4F**). On the contrary, samples belonging to other combination groups didn’t show these trends which is in line with a potentially lower immune-mediated pressure during tumor evolution.

While effective antitumor immunity hinders the accumulation of immunogenic neoepitopes in cancer samples^27^, such peptides accumulate in samples characterized by an inadequate immune response^28,29^. As metastatic samples are at a late stage of tumor evolution, we expected to find the accumulation of neoantigens in the detrimental combination group in parallel with the signs of immune evasion. Using established methods, we determined the presentation probability of each mutation by HLA-I and II molecules in each sample^29^. In line with expectation, we found an elevated number of effectively presented mutations in samples of the detrimental combination group (**Fig 4G**). In a similar analysis, we focused on nonsynonymous mutations, which are potential sources of neopeptides. We expected to find a relatively lower number of nonsynonymous mutations relative to synonymous ones in samples associated with an active immune response and the opposite trend in the ones associated with immune evasion. To this end, we calculated the ratio between observed and expected nonsynonymous mutation count (N_obs_/N_exp_) in each sample according to Rooney et al.^30^ (see Methods for details). In line with our previous findings, N_obs_/N_exp_ was elevated in metastatic samples of the detrimental combination group (**Fig 4H**).

In order to get a comprehensive picture of the biological processes that differ between samples belonging to detrimental and beneficial combination groups, we carried out a gene set enrichment analysis. As expected, primary samples in the detrimental combination group were associated with the activation of innate and adaptive immune responses (**Table 1**). The proliferation and activation of leukocytes and the dominance of antigen presentation characterized these samples. In metastatic samples of the detrimental combination group, immune responses and metabolic processes were downregulated compared to the beneficial group (**Table 1, Data S1**). At the same time, genes associated with aging, tachykinin signaling, neurogenesis and intermediate filament organization were upregulated in these samples (**Table 1**). The upregulation of aging-related genes in melanoma was recently reported to be associated with insufficient antitumor immune response and poor survival in melanoma^31^. Tachykinins promote and sustain tumor growth in multiple cancer types and also in melanoma^32^. Neurogenesis and activity of neurons are associated with a potentially immunosuppressive tumor microenvironment^33^, while the reorganization of intermediate filaments in melanoma cells correlates with tumor aggressiveness^34^.

**Table 1.** Results of gene set enrichment analysis. Significantly over- or underrepresented GO terms in the detrimental group compared to the beneficial combination group. The analysis was carried out separately for primary (left) and metastatic (right) samples. GO term classification was carried out by calculating semantic similarities between GO terms (see Methods for details). The table shows only a subset of terms. For a full list of terms, see Data S1.

| Primary - Upregulated |  | Metastasis - Downregulated |  |
| --- | --- | --- | --- |
| Parent term | Examples | Parent term | Examples |
| Cell activation | Lymphocyte activation;<br>T cell activation;<br>B cell activation;<br>Natural killer cell activation | Acute-phase response | Response to interferon-gamma;<br>Acute inflammatory response;<br>Innate immune response; |
| Regulation of immune system process | Positive regulation of immune system process;<br>Leukocyte mediated immunity;<br>Humoral immune response;<br>Complement activation;<br>Negative regulation of type 2 immune response | Aromatic amino acid family catabolic process | Protein processing;<br>Fatty acid metabolic process;<br>Peptidyl-methionine modification; |
| Immune system development | Leukocyte differentiation;<br>T cell differentiation;<br>B cell differentiation | Alpha-beta T cell activation | Alpha-beta T cell differentiation;<br>T cell activation;<br>CD4-positive, alpha-beta T cell activation; |
| Defense response | Innate immune response;<br>Natural killer cell mediated immunity;<br>Inflammatory response to antigenic stimulus;<br>Positive regulation of natural killer cell mediated cytotoxicity | Humoral immune response mediated by circulating immunoglobulin | B cell mediated immunity;<br>Humoral immune response;<br>Leukocyte mediated cytotoxicity; |
| G protein-coupled receptor signaling pathway | Immune response-regulating signaling pathway;<br>Antigen receptor-mediated signaling pathway;<br>Regulation of leukocyte mediated cytotoxicity | Metastasis - Upregulated |  |
| Cell killing | Xenobiotic metabolic process;<br>Neurotransmitter metabolic process;<br>Phagocytosis | Parent term | Examples |
| Lymphocyte migration | T cell migration;<br>Regulation of lymphocyte chemotaxis;<br>Mononuclear cell migration | Aging | Neuron differentiation;<br>Dendrite development |
| Antigen processing and presentation of peptide antigen | Peptide antigen assembly with MHC class II protein complex;<br>Antigen processing and presentation;<br>Antigen processing and presentation of exogenous peptide antigen via MHC class II | Intermediate filament-based process | Cell-cell adhesion via plasma-membrane adhesion molecules;<br>Homophilic cell adhesion via plasma membrane adhesion molecules;<br>Intermediate filament organization;<br>Synapse organization |
| Positive regulation of cytokine production | Interferon-gamma production;<br>Positive regulation of interferon-gamma production | Tachykinin receptor signaling pathway | Regulation of respiratory gaseous exchange |

Finally, it was reported that while abnormal DNA repair is associated with tumor development, proper DNA repair mechanisms are needed for tumor progression and immune evasion^35^. It has recently been suggested that elevated DNA repair exerts its pro-tumor effect through the downregulation of the STING pathway^35^. Importantly, the upregulation of DNA repair is especially common in melanoma^36–38^. The abundance of several DNA repair proteins was determined previously and collected in the TCPA database^39^. The levels of these proteins were either significantly elevated or showed a tendency of overexpression in metastatic samples belonging to the detrimental combination group (**Fig 4I**) suggesting that immune evasion in these tumors is potentially achieved through augmented DNA repair, too.

In sum, our results suggest that detrimental combinations of HLA class I and II alleles are associated with an active and effective antitumor immune response in primary samples. At the same time, strong immune selection pressure in these tumors promotes immune evasion resulting in an ineffective immune response in metastases. Importantly, the results of the survival analysis suggested that this insufficient antitumor immune response cannot be reversed with ICB therapy.

### Both beneficial and detrimental allele combinations present melanoma-associated mutations effectively

Next, we aimed to identify certain peptide-binding patterns of HLA alleles that could explain our findings. We collected 1,128,740 melanoma mutations from the COSMIC database^40^. We determined the ability of each HLA class I and II allele to present each mutation using established methods (see Methods for details). We also examined the binding properties of HLA molecules in neutral combinations as a control. First, we assumed that the similarity between the binding specificities of HLA-I and HLA-II molecules in a given combination could have a relevant effect on the immune recognition of cancer cells. To this end, we determined the relative frequency of amino acid substitutions presented by HLA-I and by HLA-II molecules in each combination. We found that HLA-I and II molecules in beneficial and detrimental combinations are specific for highly similar mutations in contrast with neutral ones (**Figure 5A**). Based on this, we expected to find a higher number of mutations presented by both HLA-I and II molecules in beneficial and detrimental combinations. These mutations could be especially immunogenic as they potentially induce both CD4+ and CD8+ T cell-mediated immune responses^8,9^. As expected, HLA-I and II molecules in beneficial and detrimental combinations were more likely to jointly present melanoma-associated mutations than the ones in neutral combinations (**Figure 5B**). Next, we aimed to understand how the presentation of melanoma-associated mutations can be associated with the opposite outcomes in the two combination groups. We examined the composition of presented amino acid substitutions in each group. Detrimental combinations were more likely to present glutamate-lysine (EK) substitutions, while beneficial ones preferred serine-phenylalanine (SF) and leucine-phenylalanine (LF) substitutions **(Figure 5C)**. All these substitutions are especially common in melanoma tumors (**Figure 5D**). Importantly, EK substitutions were classified as radical mutations by previous studies^41^, which are associated with pronounced immunogenicity. At the same time, SF and LF substitutions are nonradical ones and, thus, are potentially less likely to induce an immune response. We propose that in the case of beneficial combinations, an effective immune response develops only in metastases because these less immunogenic substitutions reach a sufficient number only in late tumor stages. This could be further augmented by the accumulation of tryptophan-phenylalanine substitutions, which occur in large numbers during the progression of many tumors due to tryptophan depletion^42^.

**Figure 5.**
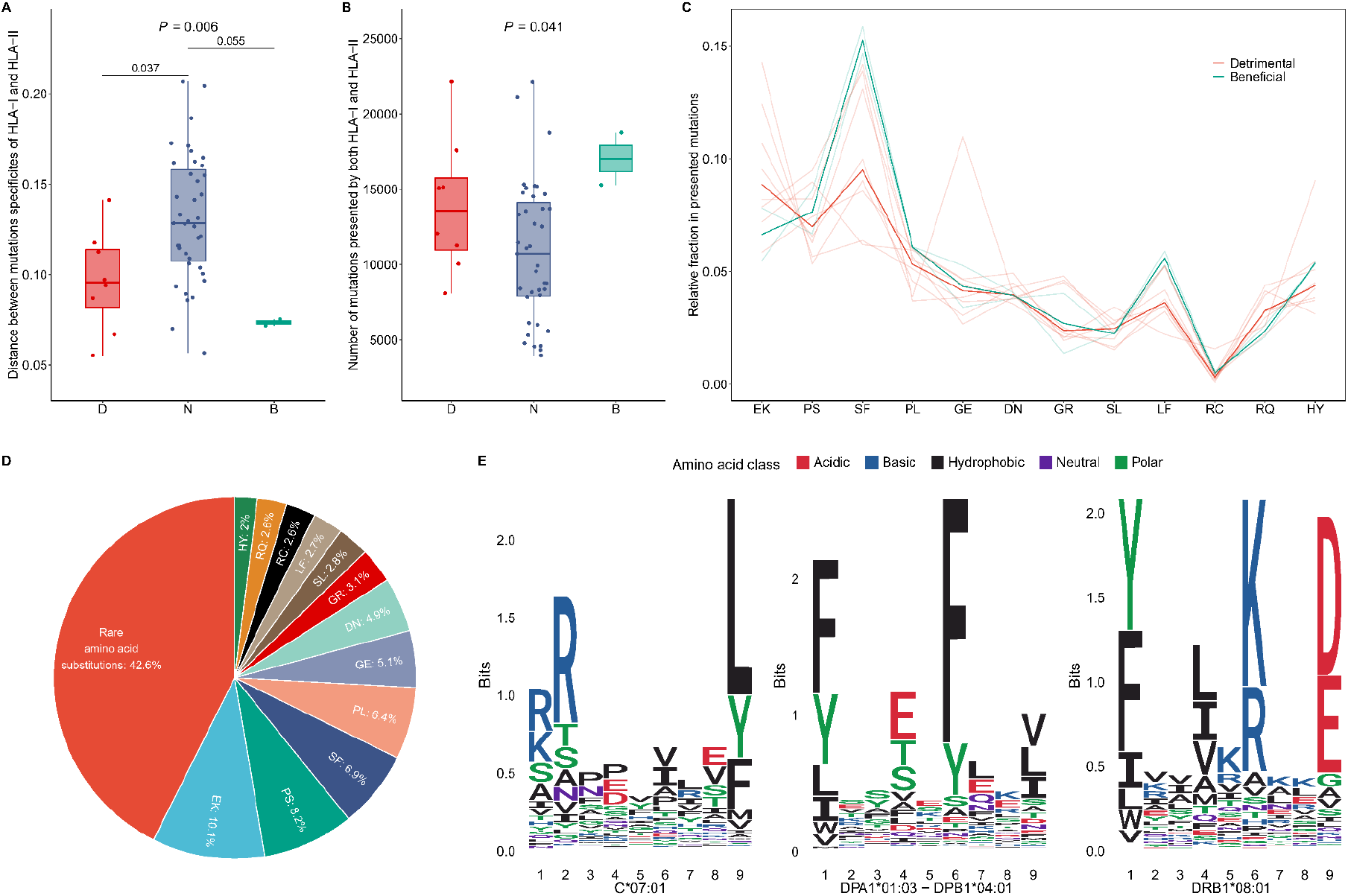
The presentation of melanoma mutations by HLA-I and II molecules. **A)** The Euclidean distance between the relative fraction of presented amino acid substitutions by HLA-I and HLA-II molecules is shown for each combination. (n = 8, 39 and 2 in the D, N and B groups, respectively) The numbers above horizontal lines indicate FDR-corrected P values of pairwise Wilcoxon’s rank sum tests. **B)** The number of mutations presented by both HLA-I and II molecules in each combination is indicated (sample counts are the same as for panel A). **C)** The parallel coordinate plot indicates the fraction of different amino acid substitutions in mutations presented by both HLA-I and II molecules in each combination. The separate combinations are shown with light colors, while group medians are shown with dark colors. Only substitutions reaching at least 2% in all melanoma mutations are shown (n = 8 and 2 in the D and B groups, respectively). **D)** The pie chart shows the fraction of amino acid substitutions in melanoma samples of the COSMIC database. Mutations not reaching 2% prevalence were classified as rare substitutions. On the top of figures A and B, two-sided P values of Kruskal-Wallis tests are shown. **E)** The sequence logos of 9-mer peptides bound by HLA-C*07:01, and the binding core region of 15-mer peptides bound by HLA-DPA1*01:03-DPB1*04:01 and DRB1*08:01 are shown (see Methods for details). The height of letters indicates the amount of information (in bits) associated with a given amino acid at the given position.

The results of binding prediction suggest that both detrimental and beneficial combinations are able to present a high number of melanoma-associated mutations effectively. At the same time, they are specific for different mutation classes. The case of HLA-C*07:01 is especially interesting, as it is included in a beneficial and a detrimental combination, too. The allele is specific for positively charged amino acids, like lysine and arginine at the first and the second position, and for the hydrophobic leucine and phenylalanine at the ninth position of nonameric peptides (Figure 5E). In the beneficial combination, its pair, HLA-DPA1*01:03-DPB1*04:01 has minuscule specificity for polar amino acids, while it is highly specific for phenylalanine. At the same time, in the detrimental combination, its pair, HLA-DRB1*08:01 is highly specific for lysine and arginine at the sixth position of the binding core. We speculate that the similarly high specificity for positively charged amino acids of both the HLA-I molecule C*07:01 and the HLA-II molecule DRB1*08:01 is responsible for the early cytotoxic immune response in primary tumors, which promotes immune evasion in later stages. On the contrary, a cytotoxic immune response develops only in metastases, when less immunogenic hydrophobic amino acids are presented by both C*07:01 and DPA1*01:03-DPB1*04:01 in the beneficial combination.

## Discussion

In our study, we examined the combined antitumor effect of HLA class I and II variants in melanoma patients. We found allele pairs that not only predicted the survival upon ICB immunotherapy but had a prognostic effect in ICB-naive melanoma patients. Importantly, these effects were prominent in late-stage patients suggesting the role of tumor-host co-evolution in our findings.

Detrimental combinations were associated with a pronounced cytotoxic immune response in primary samples exerting strong selection pressure on tumor cells, which was reflected by low ITH (**Figure 4A, Figure 6**). Strong antitumor immunity selects for clones characterized by effective immune evasion^28^. Indeed, the downregulation of HLA and the loss or downregulation of neoantigen-coding genes were prominent in metastatic samples (**Figure 4C, E and F, Figure 6**). Luksza and colleagues recently reported that in the case of the non-immunogenic cancer type, pancreatic adenocarcinoma, samples of long-term survivors were characterized by immunoediting and showed no signs of immune evasion^27^. Our results suggest that in the highly immunogenic cancer type, melanoma, immune evasion is common and has a devastating effect on patient survival (**Figures 1 and 2**). Importantly, the effect of detrimental combinations was superior to beneficial ones in samples carrying both types (**Figure S3**), which suggests that immune evasion in these samples cannot be compensated by the presentation of neopeptides by beneficial combinations (**Figure 5**). Both HLA-I and II molecules in detrimental combinations bound and presented the radical EK mutations effectively. The accumulation of EK substitutions in tumors, especially melanoma, has already been reported^43^. As glutamate (E) is negatively, while lysine (K) is positively charged, these mutations cause significant alterations in the peptide sequences and, thus, are more immunogenic^41^.

**Figure 6.**
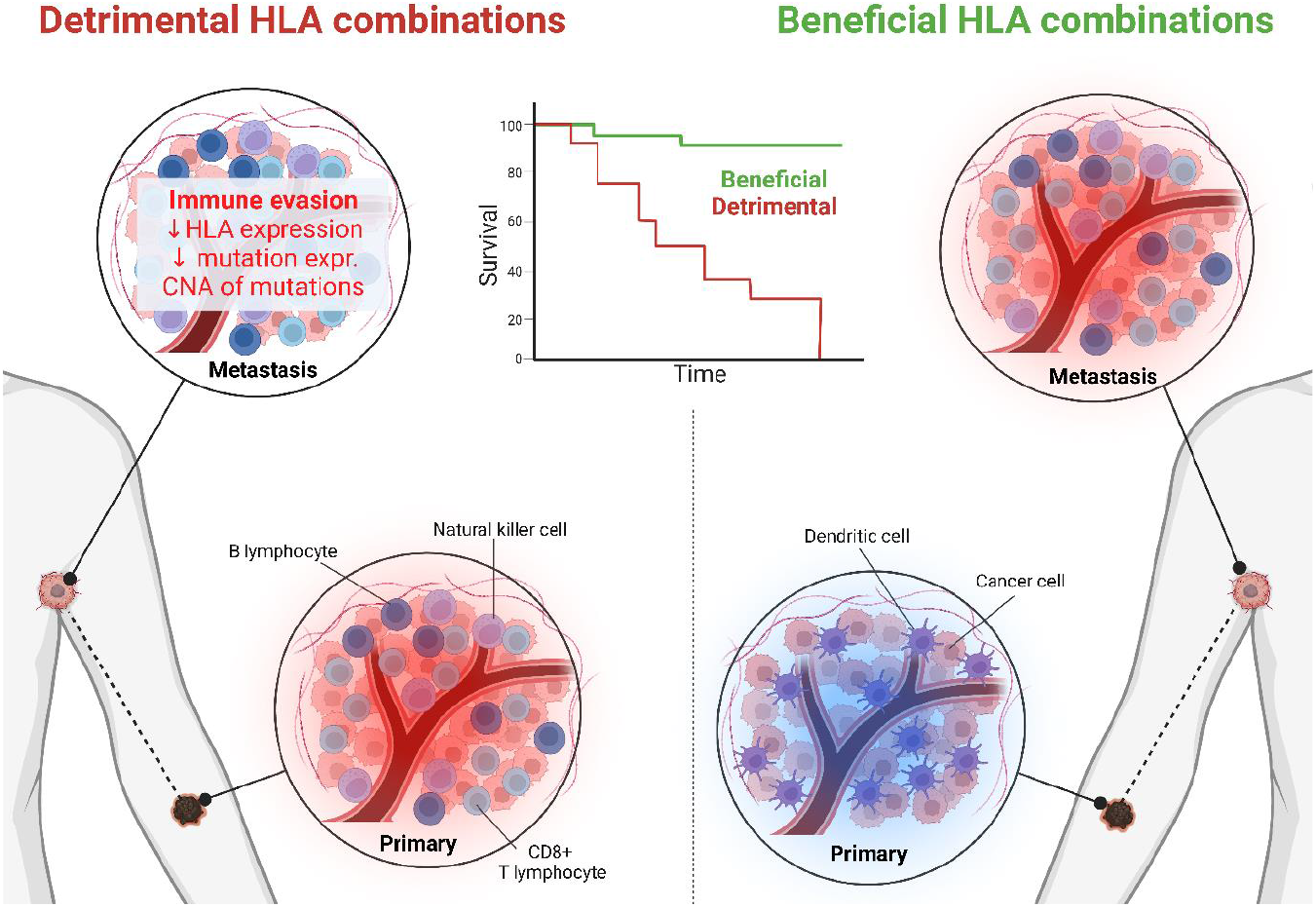
Different HLA-I and II combinations are associated with antitumor immune response in different disease stages. Detrimental combinations come with a highly active cytotoxic immune response in primary tumors leading to immune evasion in metastases. At the same time, primary tumors in patients carrying beneficial combinations are immune-cold, but metastatic samples show signs of inflammation.

Beneficial HLA combinations were associated with immunological non-responsiveness to primary melanoma samples resulting in a low number of cytotoxic T cells and a high number of dendritic cells in the TME (**Figure 3, Figure 6**). Importantly, individuals carrying these combinations were more likely to experience the recurrence of disease after the excision of the primary tumor (**Figure 2D**), which also indicates an inefficient antitumor immune response in primary tumor samples. In metastases, we found signs of effective cytotoxic immune response resulting in better survival (**Figures 1, 2 and 3**). Both HLA class I and II molecules in these combinations presented melanoma-associated neopeptides effectively (**Figure 5B**). However, they bound less immunogenic substitutions, like SF and PL (**Figure 5C**). It was recently reported that tryptophan to phenylalanine (F) substitutions also accumulate in numerous cancer types due to tryptophan depletion, which could further contribute to the positive effect of these combinations in later tumor stages^42^. The relationship between the binding specificities of beneficial combinations and the accumulation of dendritic cells in primary tumors needs further research. These cells have a multifaceted role in antitumor immunity. Our results support previous findings reporting a positive effect of dendritic cells on antitumor immunity in certain stages of tumor progression^18,44^.

Our findings suggest that certain combinations of HLA class I and II alleles can accurately and robustly predict the response to ICB therapy. Individuals carrying beneficial combinations were more likely to respond to treatment, while the ones carrying detrimental combinations had poor survival (Figure 1). It is important to highlight that individuals carrying detrimental combinations potentially have an effective antitumor immune response only at early tumor stages (Figure 3) suggesting that they could potentially gain an advantage from ICB therapy during this time period. This is in line with recent directions in therapeutic strategies proposing that ICB therapy could be used more effectively in early tumor stages or in a neoadjuvant setting because many late-stage tumors have undergone immunoediting or have an immunosuppressive TME^45–47^. At the same time, patients carrying beneficial combinations most likely respond to ICB therapy at later tumor stages as their early-stage tumors are immune-cold. Importantly, while we reported our results in melanoma, the joint effect of HLA class I and II variants in other tumor types should be investigated in future research. Similarly, the effects of strong immunological selection on tumor development should be further investigated in association with other factors that influence the strength of antitumor immunity like sex and age.

## Methods

### Identification of HLA class I and II combinations and survival analysis

We used survival, tumor mutational burden and HLA genotype data of 269 ICB-treated melanoma patients published by Chowell et al.^6^. HLA-DRB1 variants for several patients were reported as HLA-DRB1*01 by the authors. We treated them as HLA-DRB1*01:01 because these alleles are by far the most common ones in the HLA-DRB1*01 allele group (http://www.allelefrequencies.net/). Data on RECIST classification and clinical benefit were acquired from refs.^48–50^. As previously^51,52^, the long-term benefit was defined as a complete or partial response to the treatment. HED values were calculated following ref^14^. The construction of Cox proportional hazard models and the visualization of Kaplan-Meier curves were done with the survival and survminer R libraries. Only combinations occurring at least ten times were examined (homozygous alleles in a given patient were counted twice). Strong LD between HLA alleles could confound our results^53^. For example, HLA-A*01:01, B*08:01 and C*07:01 are inherited in strong LD and are members of the ancestral 8.1 haplotype, which is common in the caucasian ethnicity^54^. Importantly, the B*08:01 and C*07:01 alleles occurred with the same HLA-II allele, DPA*01:03-DPB1*04:01 in two significant combinations, which suggests a potential confounding effect of strong LD. To test whether the interaction effects are independent of each other, we constructed additional Cox models including the prevalence of other combinations as covariates:

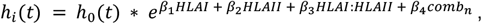

where *h_i_*(*t*) is the hazard function of individual *i, h*_0_(*t*) is the baseline hazard function; *HLAI* and *HLAII* are logical covariates indicating the presence of a given HLA-I and HLA-II molecule, respectively; *HLAI: HLAII* is the interaction term, *comb_n_* is a logical covariate indicating the presence of the n^th^ other combination with a significant effect on survival in interaction; *β*_1_, *β*_2_, *β*_4_ and *β*_3_ are regression coefficients of the three logical covariates and the interaction term, respectively. We found that the prevalence of C*07:01-DPA1*01:03-DPB1*04:01 neutralize the interaction effect between B*08:01 and DPA1*01:03-DPB1*04:01 (**Fig. S5**) suggesting that it is caused by the strong LD between C*07:01 and B*08:01 alleles. Consequently, we discarded B*08:01-DPA1*01:03-DPB1*04:01 from further analyses.

The clinical data of melanoma patients in TCGA were downloaded from cBioPortal^55^. Patients treated with ICB therapy were excluded using data on the NCI Genomic Data Commons portal^56^. Plots indicating patient history were created by using data in the data_timeline_status.txt file available on cBioPortal.

### Examining melanoma samples of TCGA

Mutational, CNA and gene expression data of melanoma samples in TCGA were downloaded from cBioPortal^55^. Data on the cellular composition of the tumor microenvironment (lymphocytes and dendritic cells) and T cell receptor diversity was acquired from ref^17^. Immune deconvolution analysis was carried out as suggested by Sturm et al^57^ using the immunedeconv R library: we used the MCP-counter algorithm for determining the levels of B cells, NK cells and the cytotoxicity score^20^ and the xCell algorithm for non-regulatory CD4+ T cells^21^. Data on ITH generated by the CloneSig software were downloaded from ref.^23^. The ratio between observed and expected missense mutation count was calculated according to Rooney et al.^30^. Briefly, we examined 192 mutational spectra, each of which was based on the original base, the mutated base, one base upstream and one base downstream of the mutated base. For each spectrum, we estimated *<u>N</u>_s_*, which is the number of non-silent mutations per silent mutations by examining all melanoma samples. Using these rates, we then calculated the number of expected non-silent mutations by considering only the mutational spectra of silent mutations in each sample. RSEM expression values of HLA and β2-microglobulin genes were downloaded from cBioPortal. CNA data of HLA and neoantigens were assessed using the data_CNA.txt file downloaded from cBioPortal. CNA status values smaller than zero were considered as losing one or both copies of HLA, B2M and neoantigen-coding genes.

### Assessing the clonality, CNA and downregulation of missense mutations

The clonality of mutations was calculated using mutation data on cBioPortal with the following formula:

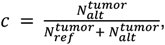

where 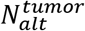 is the number of mutated, while 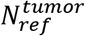 is the number of reference nucleotides detected in the tumor samples. 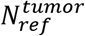 values were corrected with tumor purity using the following formula:

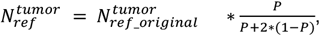

where 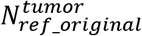 are non-corrected reference nucleotide count values while *P* is consensus tumor purity value acquired from ref.^58^. Mutations with a *c* value higher than 0.33 were considered clonal ones. We examined CNA of only subclonal mutations because we expected that the clonal or subclonal loss of the mutated gene is unlikely to result in mutations that are identified as clonal. On the other hand, we focused on the expression of only clonal mutations because we expected to identify larger differences in their case. RSEM expression values of mutated genes were acquired from cBioPortal. In both CNA and expression analysis, presented mutations were defined as having an HLA-I PHBR lower than 4 and HLA-II PHBR lower than 6, while non-presented ones were defined as having both HLA-I and HLA-II PHBR values larger than 10.

### Determining neopeptides and RHBR scores in TCGA samples

Neoantigens in each sample were determined using mutational data of melanoma samples acquired from cBioPortal. Missense mutations were selected and the sequence of the mutated peptide was downloaded from the UniProt database^59^ after matching HUGO with UniProt identifiers. Mutations were discarded if the reported original amino acid did not match the one in the protein sequence. All 8-11-mers involving the substitution were determined for the prediction of HLA-I binding and 15-mers for the prediction of HLA-II binding. The binding of each 8-11-mer to each HLA-I allele in a given sample was predicted using NetMHCPan 4.1.^60^ We used binding affinity rank values for determining binding strength. The binding of each 15-mer to each HLA-II in a given sample was predicted with NetMHCIIPan 4.0.^60^ PHBR values were calculated as previously^5,10^. Briefly, for each HLA I allele, we determined the minimal binding rank percentile value for any of the generated neopeptides by a given mutation. We then calculated the harmonic mean of allele-specific minimal binding values to determine the PHBR value of the given mutation. We involved homozygous HLA-I alleles two times in the calculation. We considered mutations with an HLA-I PHBR value smaller than 1 as strongly and under 4 as weakly presented ones in a given sample. In the case of HLA-II PHBR values, we followed the same approach. However, the multiplication of values for homozygous alleles of HLA-DP and DQ loci was slightly different. We took each combination two times if one chain-encoding gene was homozygous, and four times if both chain-encoding genes were homozygous. We considered mutations with an HLA-II PHBR value smaller than 4 as strongly and smaller than 6 as weakly presented ones. HLA genotypes of TCGA patients were acquired from ref ^5^ and ^10^.

### Gene set enrichment analysis

RSEM expression data were downloaded from cBioPortal. Gene sets for Gene Ontology “Biological Process” genes were acquired using the msigdbr v7.5.1^61^ R package. To avoid division by zero, the minimum value in the whole expression dataset (0.0028) was added to all expression values. For each gene, the mean expression value was calculated in detrimental and beneficial groups, and the log2-transformed fold change values were determined by dividing the mean expression values. Gene set enrichment analysis was carried out for each comparison using the R package fgsea v.1.18.0^62^. with the following parameters: “scoretype” = “std”; “nperm” = “1000”. Only results with FDR-corrected P values under 0.25 and a P value under 0.05 were kept. We found 212 downregulated GO terms in detrimental metastatic samples (MDD) compared to beneficial ones and 12 upregulated GO terms in detrimental metastatic samples (MDU) compared to beneficial ones and 190 upregulated GO terms in detrimental primary samples (PDU) compared to beneficial samples. For decreasing redundancy, GO term simplification was carried out by “visRDAGBP” function from GOxploreR package version 1.2.6^63^. After simplification, MDD, MDU and PDU group contained 166, 11 and 147 GO terms respectively. Next, a semantic similarity matrix was calculated by “calculateSimMatrix” function (ont = “BP”, method = “Wang”) of the GOSemSim package v.2.0.0^64^. GO term clustering was carried out by “reduceSimMatrix” function, using log2 transformed GSEA P-values as “scores” and a threshold of 0.9.

### Determining the HLA-presentation of melanoma mutations in the COSMIC database

Mutations in melanoma samples were downloaded from the COSMIC database website^40^ (v96_38_ MUTANT.csv file accessed on 29/07/2022). Missense mutations were selected and the original and mutated protein sequence was determined using the UniProt ID and sequence data on the Uniprot database^59^ (accessed on 29/07/2022). Mutations were discarded if the reported original amino acid did not match the one in the downloaded sequences. The HLA-binding strength of neopeptides generated from the remaining 1,128,740 mutations was predicted using the same methods as for mutations in TCGA samples. As previously^5,10^, the lowest rank EL score value (reflecting strongest binding) of any 8-12-mer and 15-mer was determined for each mutation as a proxy of binding strength to HLA-I and HLA-II molecules, respectively. Mutations having a minimal value under 0.5 and 1 were considered bound ones by HLA-I and HLA-II molecules, respectively. We used cutoffs of strong binding to minimize false positive hits. Neutral combinations were defined as the ones having especially low and non-significant interaction term coefficients (i. e. the absolute value of the coefficient was lower than 0.2). Binding logos of HLA molecules were generated with ggseqlogo^65^. Nine amino acid-long peptides bound by HLA-C*07:01 were acquired from a large immunopeptidomics study published by Sarkizova et al.^66^s. The sequence logos of HLA-II alleles were created by selecting the 1% most strongly bound 15-mer peptides by the given allele in the COSMIC dataset and collecting the binding cores as reported by the NetMHCIIpan 4.0 algorithm.

## Acknowledgments

We thank Hannah Carter, J. Font-Burgada and Rachel Marty for sharing the HLA-I and II genotype data of TCGA patients. The results here are in part based on data generated by the TCGA Research Network (https://www.cancer.gov/tcga). The work was supported by the European Union’s Horizon 2020 research and innovation program grant No. 739593 (M.M., B.K, G.M.B); This work was supported by OTKA grant FK-142312 (M.M). This research work was conducted with the support of the National Academy of Scientist Education under the sponsorship of the Hungarian Ministry of Innovation and Technology (FEIF/646-4/2021-ITM_SZERZ). The work is supported by the ÚNKP-22-2 and ÚNKP-22-4 New National Excellence Program of the Ministry for Culture and Innovation from the source of the National Research, Development and Innovation Fund (A.T.F., G.M.B.). Figure 6. Was created with BioRender.com.

## Authorship contributions

Conceptualization: M.M.; Methodology: M.M., B.T.P., A.T.F., G.M.B., B.K., D.S.; Analysis: M.M., B.T.P., A.T.F., G.M.B., B.K., D.S.; Resources were provided by M.M.; Data curation was undertaken by M.M., B.K., G.M.B. and B.T.P.; Writing of the original draft was done by M.M.; Project administration and supervision were carried out by M.M.; Funding acquisition was the responsibility of M.M.

## Supplementary material

**Figure S1.**
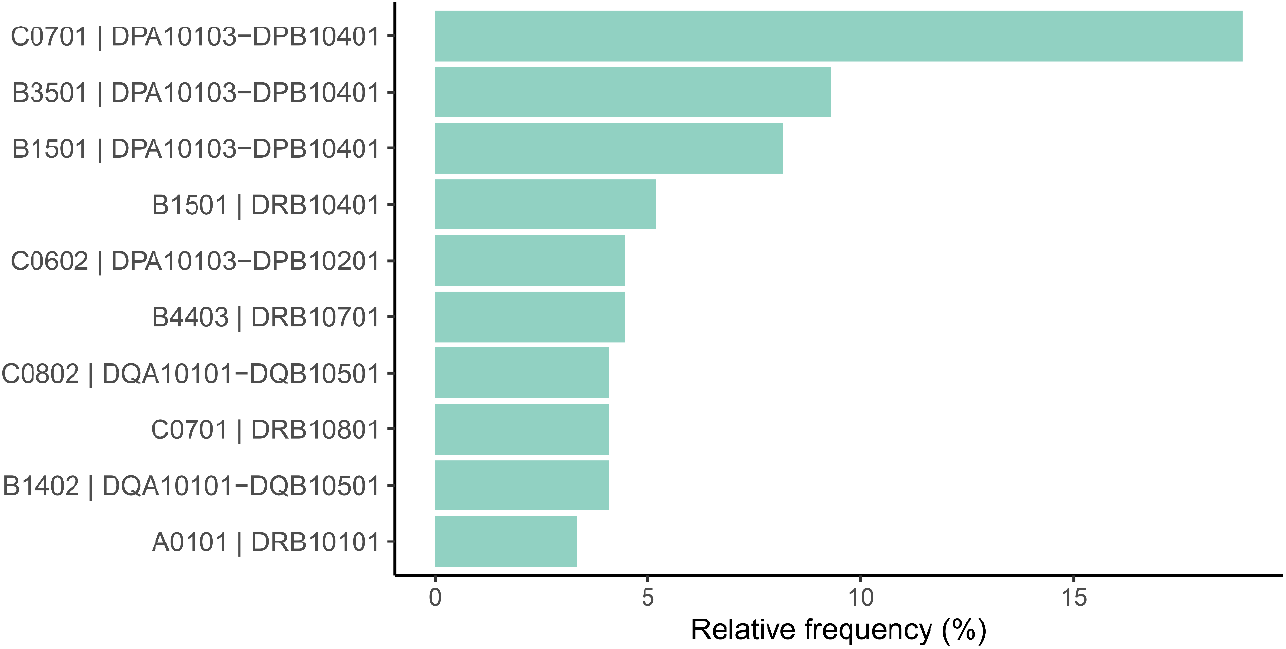
Prevalence of detrimental and beneficial combinations in the ICB-treated patient cohort. The bar chart shows the frequency of patients carrying a given combination.

**Figure S2.**
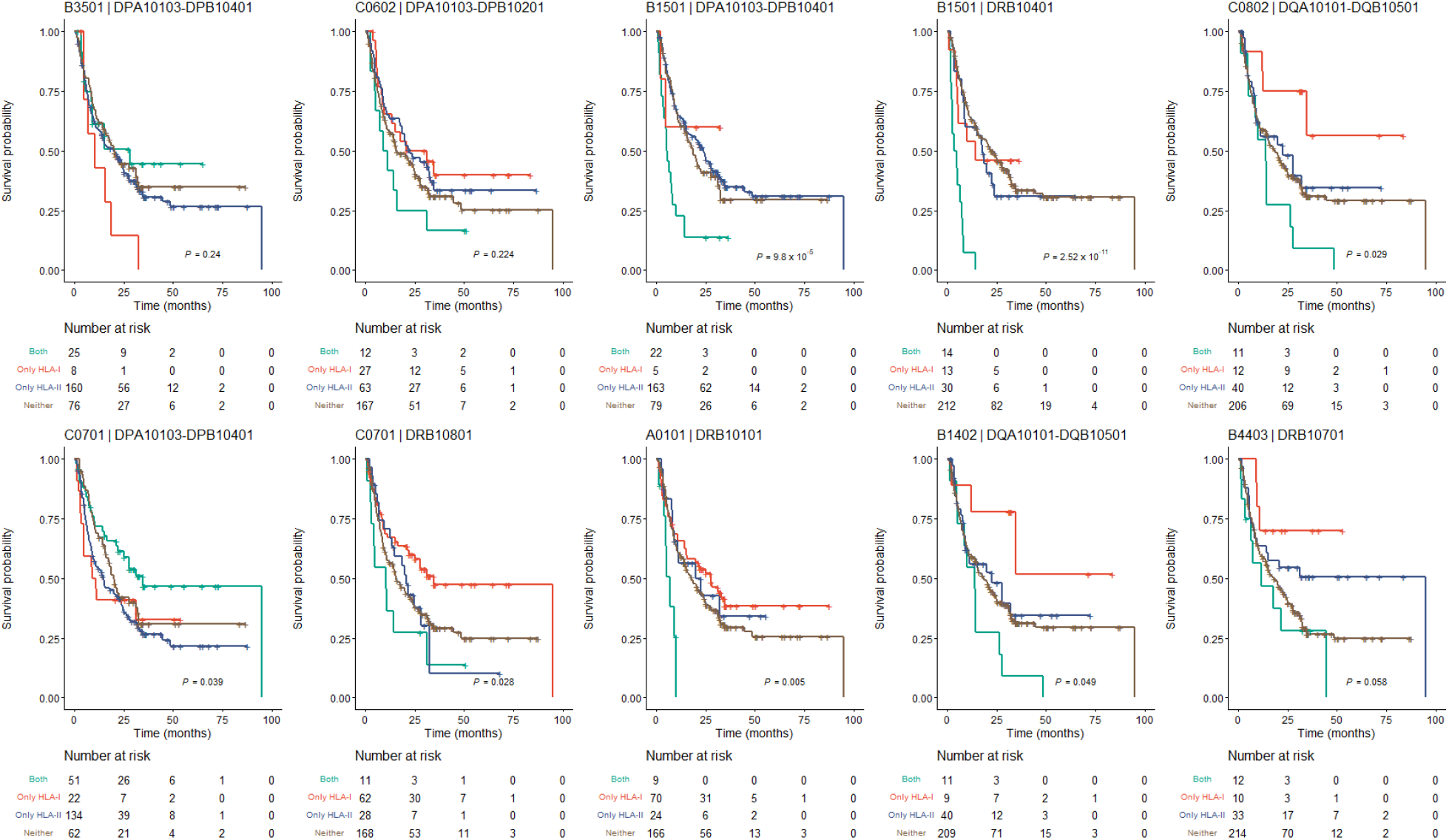
Survival of ICB-treated patients carrying different HLA combinations. Patients were grouped based on whether they carried only the HLA-I or the HLA-II molecule of the combination, both, or neither of them. Two-sided log-rank test P values are shown. Vertical axes indicate the probability of patient survival.

**Figure S3.**
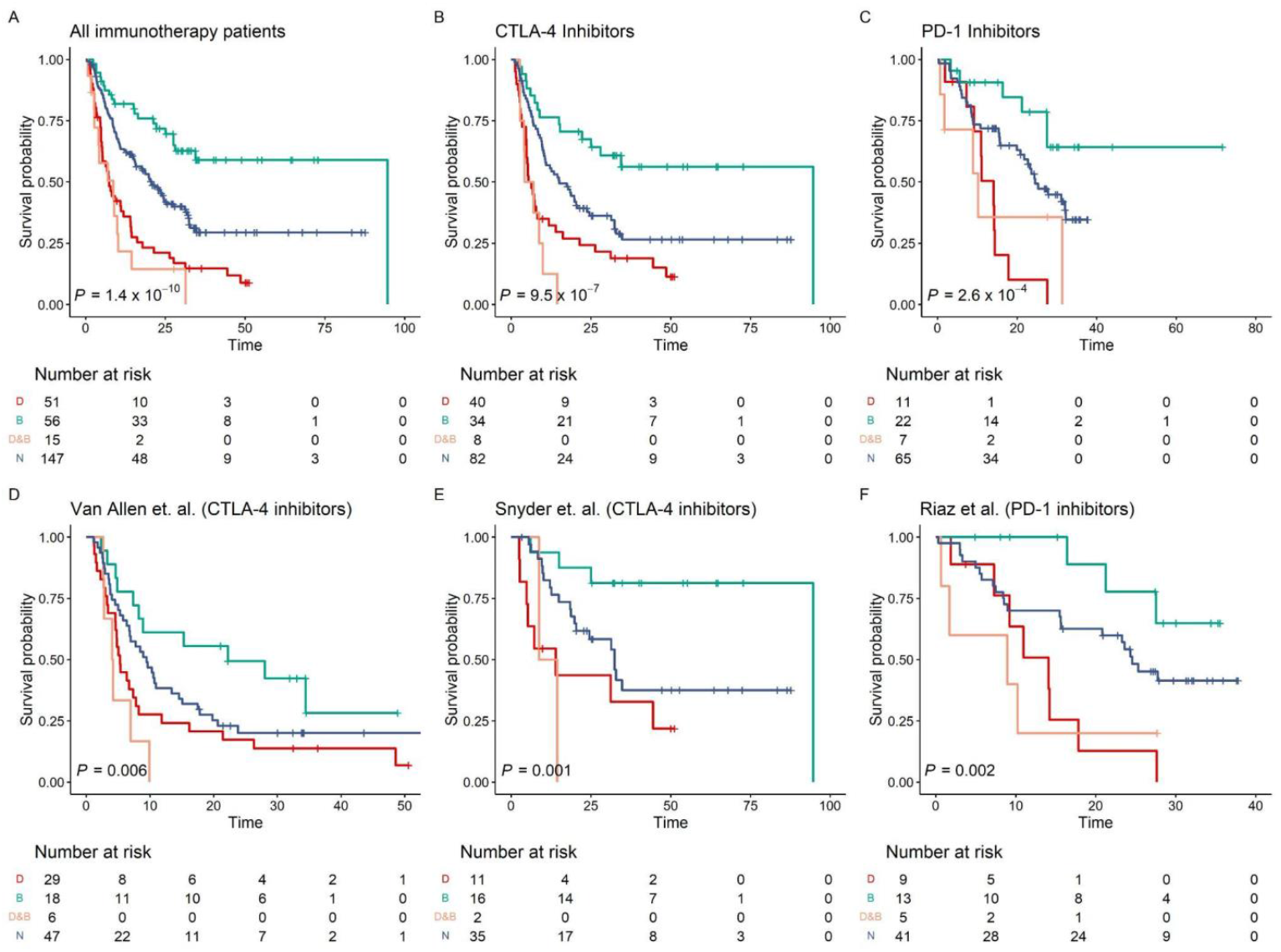
Survival of cancer immunotherapy patients carrying detrimental (D), beneficial (B), both (B&D) combinations, or only neutral ones (N). Two-sided log-rank test P values are shown. Vertical axes indicate the probability of patient survival.

**Figure S4.**
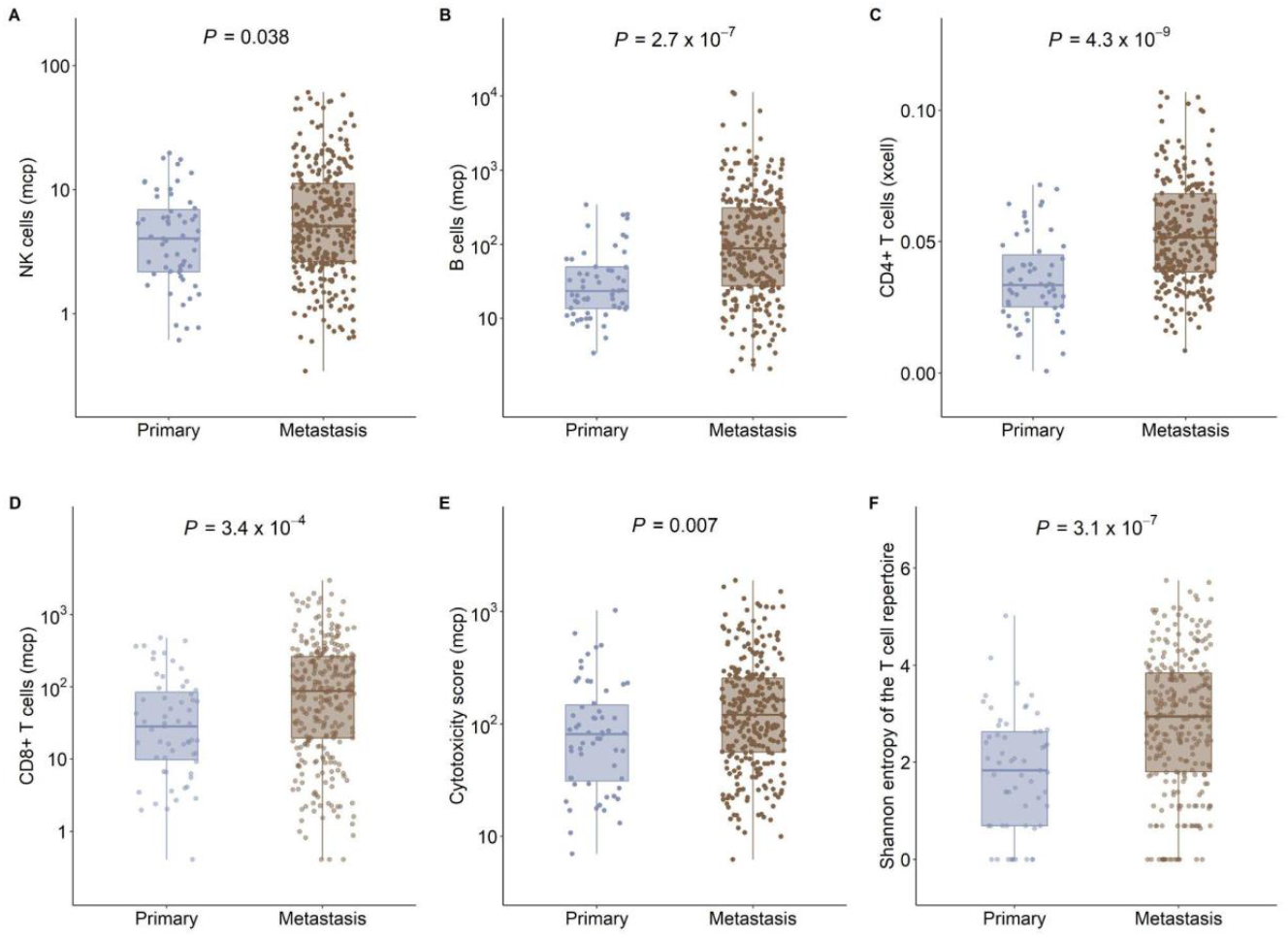
The tumor immune microenvironment in primary and metastatic samples. **A-B)** The abundance of NK cells and B cells was determined with the MCP-counter algorithm^66^ (10.1186/s13059-016-1070-5) (n = 59 and 288 in primary and metastatic tumor samples, respectively). **C)** The relative abundance of CD4+ T cells determined with the xCell algorithm^21^ (10.1186/s13059-017-1349-1) (sample counts are the same as for panels A-B). **D)** The abundance of CD8+ T cells determined with the MCP-counter algorithm^20^ (10.1186/s13059-016-1070-5) (sample counts are the same as for panels A-B). **E)** The cytotoxicity scores in samples as calculated by the MCP-counter algorithm^20^ (10.1186/s13059-016-1070-5) (sample counts are the same as for panels A-B). **F)** The Shannon entropy of the T cells in tumor samples repertoire as reported by ref.^17^ 10.1016/j.immuni.2018.03.023 (n = 59 and 269 in primary and metastatic tumor samples, respectively). On panels A to F, P-values of Wilcoxon’s rank-sum tests are indicated (on top). The bottom and top of the boxes indicate the first and third quartile, horizontal lines indicate the median, and vertical lines indicate the first quartile - 1.5*IQR and the third quartile + 1.5*IQR.

**Figure S5.**
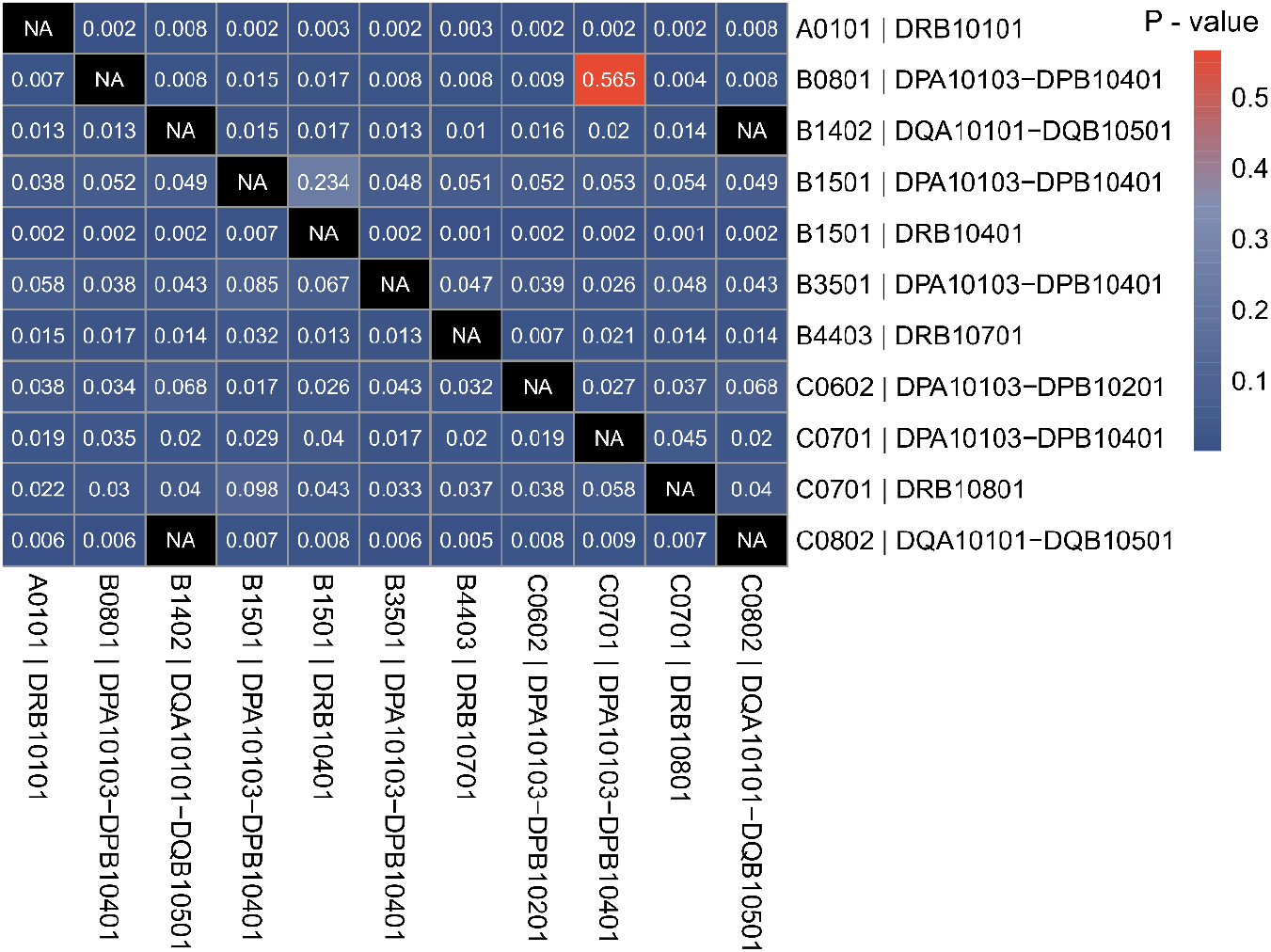
Controlling for LD in Cox survival models. The heatmap shows the P values of interaction terms between HLA-I and II alleles (in rows), when the presence of other combinations (in columns) is also included in the models (see Methods for details).

**Table S1.**
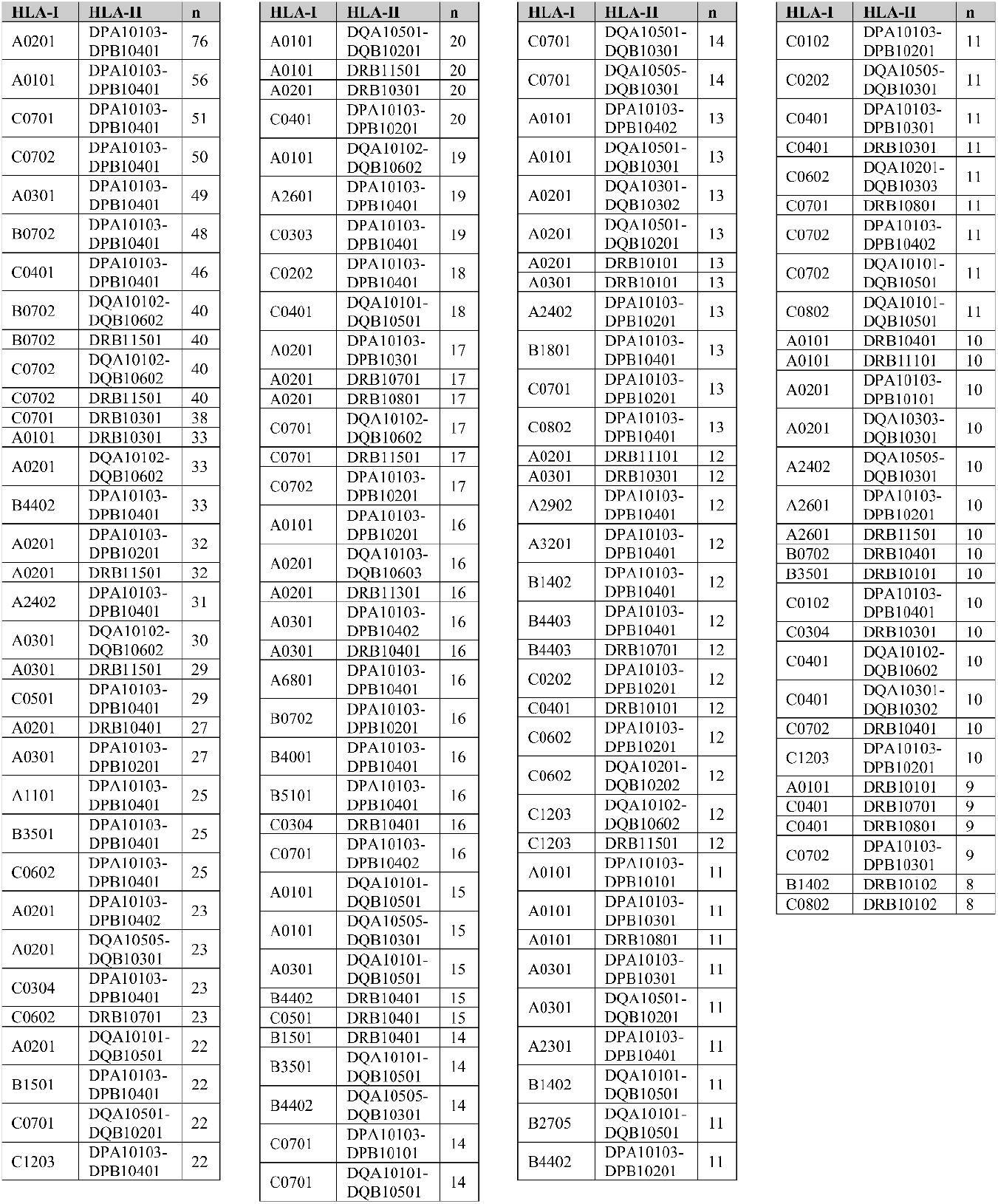
Most frequent HLA combinations in patients.

**Table S2.** Details of models with significant interaction terms. In the 5th, 7th and 9th columns, two-sided P values of z-statistics are shown. HR: hazard ratio

| HLA-I | HLA-II | Combination type | HR (HLA-I) | P-value (HLA-I) | HR (HLA-II) | P-value (HLA-II) | HR (interaction) | P-value (interaction) | P-value (Log-rank test) |
| --- | --- | --- | --- | --- | --- | --- | --- | --- | --- |
| C0701 | DPA10103<br>DPB10401 | Beneficial | 1.3 | 0.39 | 1.3 | 0.19 | 0.41 | **0.021** | **0.036** |
| B3501 | DPA10103<br>DPB10401 | Beneficial | 2.1 | 0.064 | 1.1 | 0.5 | 0.37 | **0.049** | 0.36 |
| B1501 | DPA10103<br>DPB10401 | Detrimental | 0.65 | 0.55 | 0.89 | 0.5 | 4.6 | **0.045** | 0.085 |
| B1501 | DRB10401 | Detrimental | 1 | 0.98 | 1.2 | 0.42 | 5.3 | **0.002** | 0.36 |
| B4403 | DRB10701 | Detrimental | 0.37 | 0.085 | 0.62 | 0.067 | 5.9 | **0.013** | 0.25 |
| B1402 | DQA10101<br>DQB10501 | Detrimental | 0.36 | 0.078 | 0.91 | 0.69 | 5.6 | **0.013** | 0.4 |
| C0602 | DPA10103<br>DPB10201 | Detrimental | 0.71 | 0.22 | 0.81 | 0.26 | 2.5 | **0.04** | 0.93 |
| C0802 | DQA10101<br>DQB10501 | Detrimental | 0.35 | **0.041** | 0.9 | 0.65 | 5.7 | **0.006** | 0.4 |
| A0101 | DRB10101 | Detrimental | 0.77 | 0.17 | 0.87 | 0.64 | 4.7 | **0.003** | 0.27 |
| C0701 | DRB10801 | Detrimental | 0.6 | **0.015** | 1 | 0.98 | 2.6 | **0.033** | 0.087 |

**Table S3.** Summary of Cox models for separate HLA-I and HLA-II alleles. Two-sided P values of z-statistics are shown. HR: hazard ratio

| Allele | HR | P | Allele | HR | P | Allele | HR | P | Allele | HR | P |
| --- | --- | --- | --- | --- | --- | --- | --- | --- | --- | --- | --- |
| A0101 | 0.9 | 0.5 | B0702 | 1 | 0.3 | B4402 | 0.8 | 0.3 | C0602 | 1 | 0.8 |
| A0201 | 1 | 0.6 | B0801 | 0.9 | 0.5 | B4403 | 0.9 | 0.6 | C0701 | 0.7 | 0.06 |
| A0301 | 1 | 0.6 | B1302 | 1 | 1 | B5101 | 0.9 | 0.6 | C0702 | 1 | 0.8 |
| A1101 | 1 | 0.5 | B1402 | 1 | 1 | B5701 | 1 | 0.6 | C0802 | 0.9 | 0.6 |
| A2301 | 0.6 | 0.1 | B1501 | 2 | 6.9x10<sup>-4</sup> | C0102 | 1 | 0.6 | C1203 | 0.8 | 0.5 |
| A2402 | 0.9 | 0.5 | B1801 | 0.8 | 0.3 | C0202 | 1 | 0.7 | C1502 | 1 | 0.8 |
| A2601 | 1 | 1 | B2705 | 1 | 0.4 | C0303 | 1 | 0.1 | C1601 | 2 | 0.06 |
| A2902 | 1 | 0.8 | B3501 | 1 | 0.9 | C0304 | 1 | 0.4 |  |  |  |
| A3201 | 0.6 | 0.2 | B3801 | 0.9 | 0.7 | C0401 | 0.9 | 0.8 |  |  |  |
| A6801 | 1 | 0.2 | B4001 | 0.9 | 0.8 | C0501 | 0.8 | 0.2 |  |  |  |
| DPA10103<br>DPB10101 | 0.9 | 0.6 | DQA10102<br>DQB10602 | 1 | 0.9 | DQA10501<br>DQB10201 | 1 | 0.8 | DRB10901 | 1 | 0.8 |
| DPA10103<br>DPB10201 | 0.9 | 0.7 | DQA10102<br>DQB10604 | 0.7 | 0.3 | DQA10501<br>DQB10301 | 1 | 0.9 | DRB11101 | 1 | 0.4 |
| DPA10103<br>DPB10301 | 1 | 0.5 | DQA10103<br>DQB10603 | 0.8 | 0.4 | DQA10505<br>DQB10201 | 1 | 0.9 | DRB11104 | 0.5 | 0.08 |
| DPA10103<br>DPB10401 | 1 | 0.9 | DQA10104<br>DQB10503 | 1 | 0.8 | DQA10505<br>DQB10301 | 1 | 0.9 | DRB11301 | 0.8 | 0.3 |
| DPA10103<br>DPB10402 | 1 | 0.1 | DQA10201<br>DQB10202 | 0.6 | 0.1 | DRB10101 | 1 | 1 | DRB11302 | 0.7 | 0.2 |
| DPA10201<br>DPB10401 | 1 | 0.7 | DQA10201<br>DQB10301 | 0.7 | 0.4 | DRB10102 | 1 | 0.7 | DRB11454 | 1 | 0.9 |
| DQA10101<br>DQB10301 | 1 | 0.7 | DQA10201<br>DQB10303 | 1 | 1 | DRB10301 | 0.8 | 0.4 | DRB11501 | 1 | 0.7 |
| DQA10101<br>DQB10501 | 1 | 0.5 | DQA10301<br>DQB10301 | 2 | 0.1 | DRB10401 | 2 | 0.002 | DRB11601 | 1 | 0.9 |
| DQA10101<br>DQB10602 | 1 | 0.9 | DQA10301<br>DQB10302 | 1 | 0.8 | DRB10701 | 0.8 | 0.3 |  |  |  |
| DQA10102<br>DQB10501 | 1 | 0.5 | DQA10303<br>DQB10301 | 1 | 0.2 | DRB10801 | 1 | 0.2 |  |  |  |

